# CCGM: a Compound Coarse Grain Model representation for enhanced chemotype exploration, annotation and screening

**DOI:** 10.1101/2024.12.16.628696

**Authors:** Navriti Sahni, Marcel Patek, Rayees Rahman, Balaguru Ravikumar

## Abstract

Structurally similar compounds often exhibit similar bioactivity, making similarity estimation an essential step in many cheminformatics workflows. Traditionally, compound similarity has been evaluated using diverse molecular representations, such as molecular fingerprints, compound 3D structural features, and physicochemical properties. These methods have proven effective, particularly during the early stages of drug discovery, where the primary goal is to identify initial hits from large compound libraries. However, these representation and methods often fall short during the hit-to-lead development phase, where modifications to the core scaffold or chemotype are performed and evaluated. To address this limitation, we developed the Compound-Coarse-Grain-Model (CCGM), a framework that represents structural features of a compound as nodes and edges within a simplified graph. This approach augments the pharmacophore and chemotype features of the compound within the graph, enabling the identification of compounds with similar chemotype and pharmacophore features more effectively than conventional methods. CCGM is particularly useful for when screening large libraries to identify compounds with similar chemotypes and for filtering generative designs to retain designs with similar pharmacophore features.

## Introduction

In early-stage drug discovery, potential lead candidates are typically identified and optimized through high-throughput screening (HTS) or virtual screening efforts. These processes facilitate the identification of active molecules, known as “hits”, that interacts with specific biological targets. Following initial identification, hits undergo hit expansion and hit-to-lead optimization steps, where the chemical structure is further refined to improve potency, selectivity, and drug-like properties. Identifying bioactive chemotypes— distinct chemical structures with expected biological response—is essential for the expansion and optimization phases of drug discovery [1]. Chemotype identification not only informs structure-activity relationship (SAR) analysis but also plays a pivotal role in diversifying and refining lead compounds for drug development. Furthermore, the scaffolds underlying these chemotypes form the foundational structure that enables medicinal chemistry design and optimization. They serve as the structural basis for generative designs and are critical for analyzing and comparing bioactive compounds and their analogs in the pursuit of new active molecules. To enable such searches computationally, a myriad of molecular representations has been developed for computer-aided drug design (CADD) [2]. These representations are essential for associating a compounds’ structure and molecular properties with its biological activity or other experimental endpoints [3]. For instance, structural attributes such as atom types, bond types and bond order can be represented as nodes and edges of a graph, which are then linked to observed physicochemical property or biological activity to train machine-learning and deep-learning models [4].

Traditional approaches, like compound fingerprints, such as extended-connectivity fingerprints (ECFP), have proven valuable in similarity searches for identifying and categorizing bioactive compounds [5]. However, these methods often fail to capture embedded chemotype information, which is critical for understanding specific protein-ligand interactions and follow-up optimization. Similarly, 3D representation of compounds has been instrumental in virtual screening studies, particularly in docking. While effective, these methods are often computationally intensive, limiting their scalability [6]. Beyond molecular representations, recent advancements in scoring functions and similarity search algorithms have significantly accelerated the virtual screening process. Notable examples include ligand-centric pharmacophore screening approaches such as ROCS, DeCAF and Roshambo [7–9]. These methods often integrate molecular alignment methods with GPU-acceleration services to facilitate expedited virtual screening tasks. However, they may filter out compounds with unconventional chemotypes, thereby limiting the exploratory scope. While these approaches categorically estimate shape and pharmacophore similarities, they often fail to distinguish the core chemotype of a compound from its R-groups an essential consideration in hit-2-lead optimization studies. Addressing the limitations associated with molecular representations and similarity search methods is critical for advancing drug discovery and improving the precision of virtual screening workflows. Integrating geometric features, such as exit vectors their positions and angles, incorporating atoms physicochemical attributes for chemotype exploration, distinguishing core scaffold and R-groups for structure-activity relationship (SAR) studies, are vital for enhance various facets of drug development. These functionalities can improve key process, including virtual screening, medicinal chemistry design, and curating designs from generative models.

To address these unmet needs, we developed Compound Coarse Grain Model (CCGM), a framework that employs a reduced graph representation of a compound to enable similarity searches. In CCGM, complex molecular features of a compound such as ring systems, linkers, and terminal atoms are abstracted into nodes, while bonds represented as edges. This reduced representation allows CCGM to embed key molecular features, structural properties and pharmacophore characteristics into the graph’s nodes and edges. In turn, this enables optimal molecular alignment and facilitates the estimation of similarity scores that effectively combines chemotype and pharmacophore similarities. Furthermore, the incorporation of node and edge weights in CCGM allows users to distinguish between core chemotypes and R-groups, providing greater insights during SAR studies. To validate our approach, we benchmarked its performance by evaluating its ability to screen both similar and diverse chemotypes, comparing the results against existing similarity search methods. We further demonstrate the utility of CCGM in screening large chemical libraries and its effectiveness in filtering structure-relevant compounds from generative models. We believe that CCGM can become an integral component of a computational and medicinal chemist’s toolkit, addressing critical requirements for virtual screening in drug discovery.

### Implementation

The Compound Coarse Grain Model (CCGM) employs a reduced graph representation of a compound. This representation allows users to distinguish a compound’s core chemotype from its decoration groups (R-groups). This approach transforms molecular features such as rings, terminal atoms, and linkers into nodes and edges, with all the rings atoms reduced to a single node. Pharmacophore properties of the molecule are incorporated into the reduced graphs as attributes. CCGM allows users to define the core chemotype of a compound for similarity searches by adjusting the weights to nodes and edges to the scaffold of interest. A schema illustrating this transformation of a compound to its reduced weighted graph with node and edge attributes is depicted in Fig. 1. This representation is crucial for cheminformatics workflows, as hits progress from discovery to the optimization phase in drug discovery, where the objective is to refine and enrich the chemotype for desired properties. The process begins by identifying a promising compound with a specific scaffold or chemotype (hits) from early screening studies and refining these hits to achieve the desired biochemical and physicochemical properties. In such cases, a scaffold-based similarity metric like CCGM is highly valuable, as it enables the identification of compounds with similar scaffolds and pharmacophore properties, thereby effectively guiding the design of lead molecules. The CCGM module is implemented as a python package with all the required dependencies installed through the provided conda environment. There are dedicated classes in package that enable parallelization of the similarity calculations across multiple CPU processors, enhancing computational efficiency. The CCGM module is written in Python and is dependent on several freely available python packages such as RDKit , NetworkX [10] and datamol.

**Figure 1.**
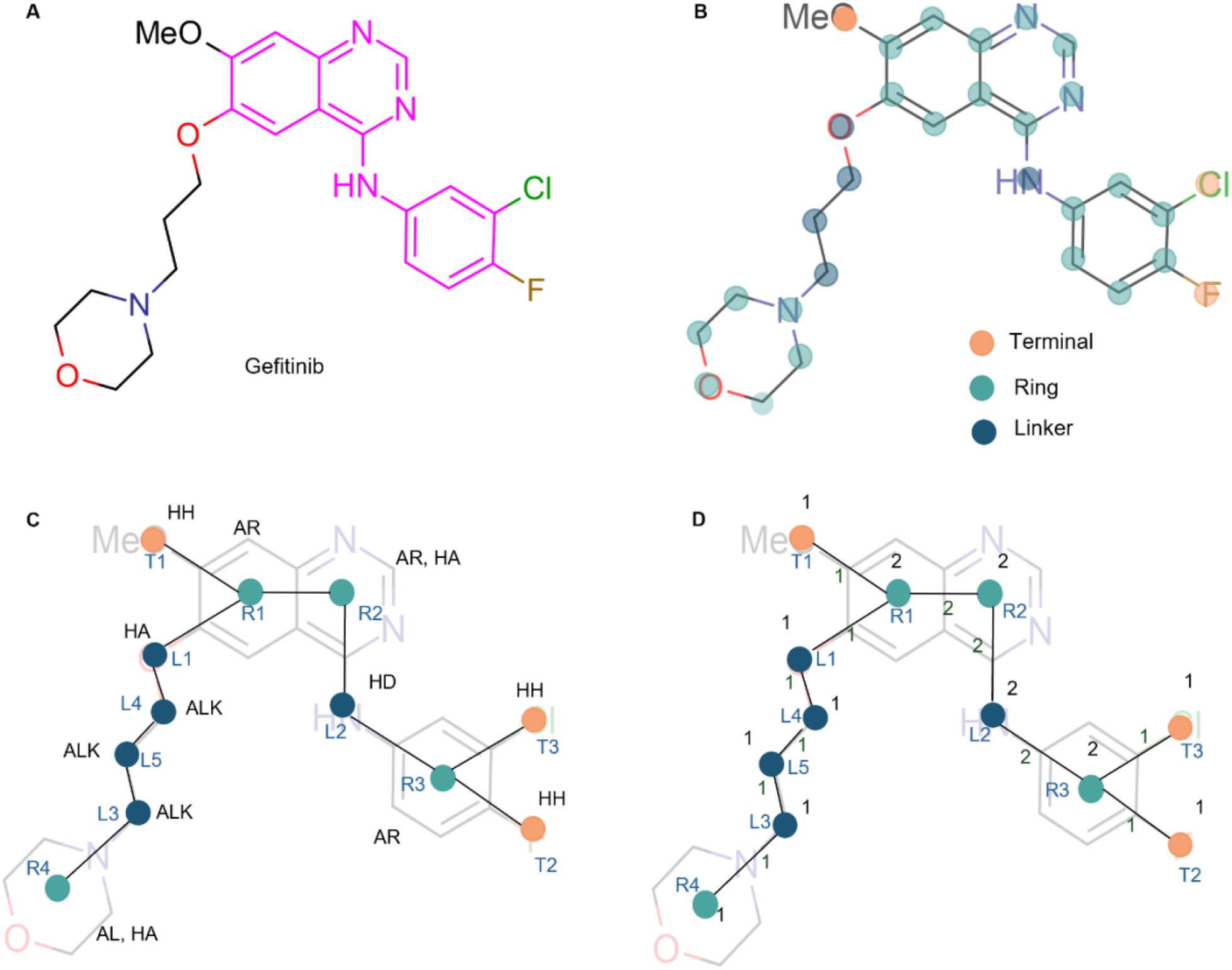
A schematic illustration of CCGM representation and accompanying attributes: (A) A 2D representation of gefitinib oriented using Ketcher, the core scaffold is highlighted in pink. (B) Molecular structures are transformed into nodes and edges. Node classification categorizes nodes as terminal (T), linker (L), and ring center (R). (C) Each node is annotated with pharmacophore features. (D) Node and edge weights of the reduced graph that enable weight CCGM similarity calculations.

#### Chemotype definition and alignment

Once an initial scaffold is selected, users need to submit the corresponding template molecule containing the scaffold in Structure Data File (.*sdf*) format. The 2D coordinates of the compound should be oriented as desired, using standard molecular editing software, such as Ketcher [11]. The list of query molecules (library) should be provided as a .*csv* file, including the molecules Simplified Molecular Input Line Entry System (SMILES) notation [12] and their corresponding IDs. Each SMILES string is converted into its respective molecule object to calculate similarity with the template molecule. To eliminate redundant node assignments, specific terminal functional groups in both the template and query molecules are abbreviated. This condense representation of the compounds are used for alignment, the CondenseMolAbbreviations module provided by RDKit is used for this step. Additionally, to ensure bond lengths are comparable across molecules, the bond distances are first normalized. The Maximum Common Substructure (MCS) shared between the query and template molecule are identified using the rdFMCS module [13], which is then used to align the query and template molecules, as illustrated in Fig. 2A.

**Figure 2.**
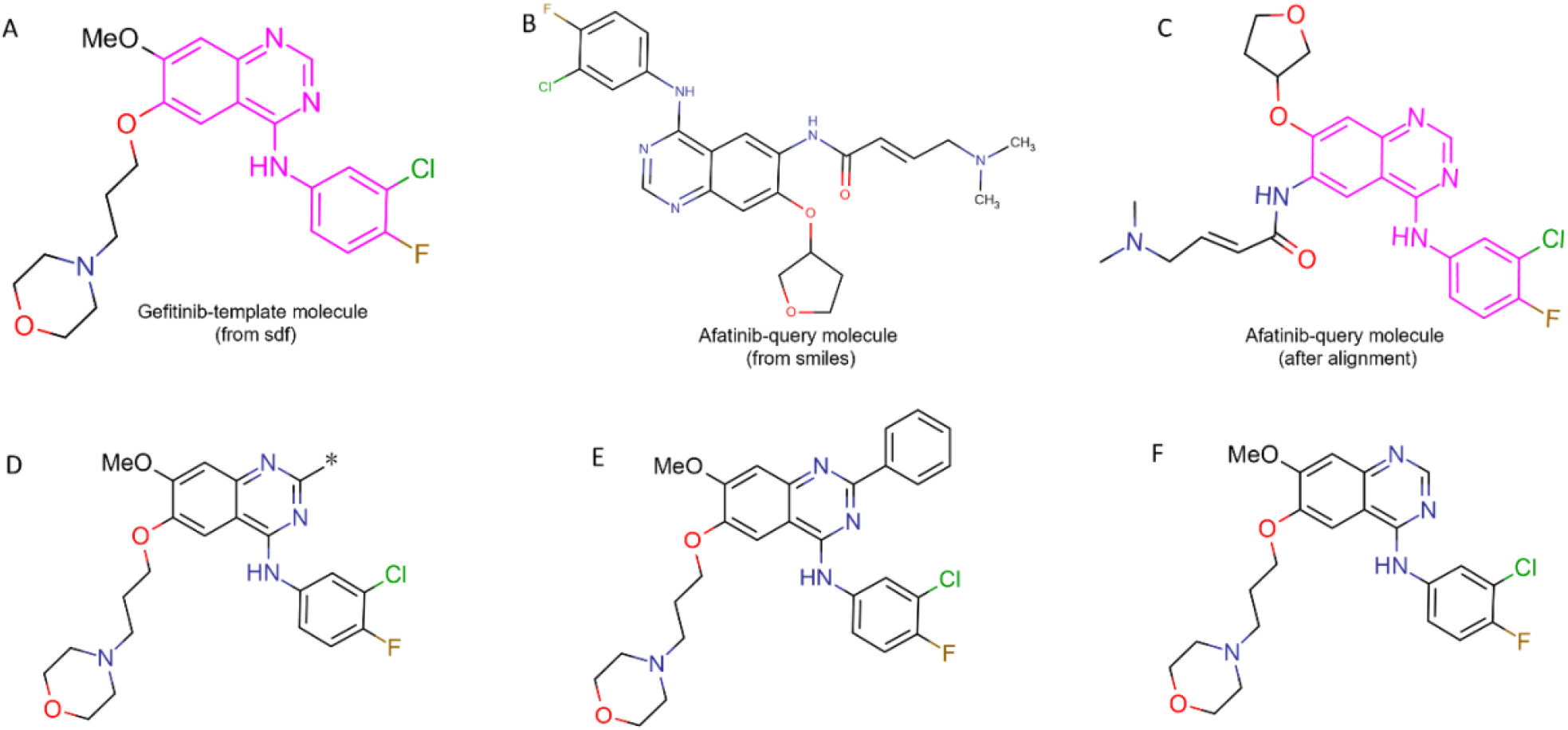
(A) An example where Gefitinib is used as the template molecules and afatinib as the query compound to estimate CCGM similarity, the user defined core scaffold is highlighted in pink. (B) The 2D coordinates for afatinib are obtained from the SMILES representation of the molecule. (C) Aligned query molecule afatinib based on the maximum common scaffold (MCS), shared between the template and query molecules. (D) A dummy atom (*) which is used to represent an exclusion area, defining areas where negative weights are applied for that exclusion zone. (E) Impact of negative weights on CCGM scores: Assigning a negative weight of (for example -10) to a dummy atom causes the addition of a phenyl ring node, replacing the dummy atom or exclusion area, would significantly lower the CCGM score (from 1 with default weights of 1 to 0.24). (F) When no substitution is made in place of the dummy atom or exclusion area, assigning a negative weight of -10 results would result in an increase in the CCGM score (from 0.95 with default weights of 1 to 1.7).

#### Graph representation of molecules

Once aligned, both the template and query compounds are converted into their respective graph representations. To facilitate this conversion, atoms in each molecule are categorized as ring, terminal, or linker atoms, in certain cases, fused ring atoms are assigned a specific category. Along with these categories, the indices of their 2D coordinates, and ring centers (for ring atoms) are stored. This data is then summarized to create a nodes lookup table and an adjacency matrix, which captures the node and edge interaction. Using the Network module, this information is used generate the reduced graph representation, that forms the backbone of the CCGM. The alignment quality between the template and query molecules are further verified using their respective graphs and their MCS.

#### Augmenting pharmacophore features

To provide a more comprehensive representation of a molecule’s bioactive potential and to enable pharmacophore similarity searches, CCGM incorporates a pharmacophore lookup table. Graph nodes are enriched with pharmacophore features essential for understanding the molecule’s biological activity, as shown in Fig. 1B. Like DeCAF, specific pharmacophore annotations such as: hydrogen bond acceptors (HA), hydrogen bond donors (HD), hydrophobic regions (HH), internal hydrogen bonds (INT), basic amines (BA), and aromatic ring systems (AR), are assigned to each nodes as attributes. A detailed annotation of these pharmacophore features is provided in Table 1.

**Table 1.**
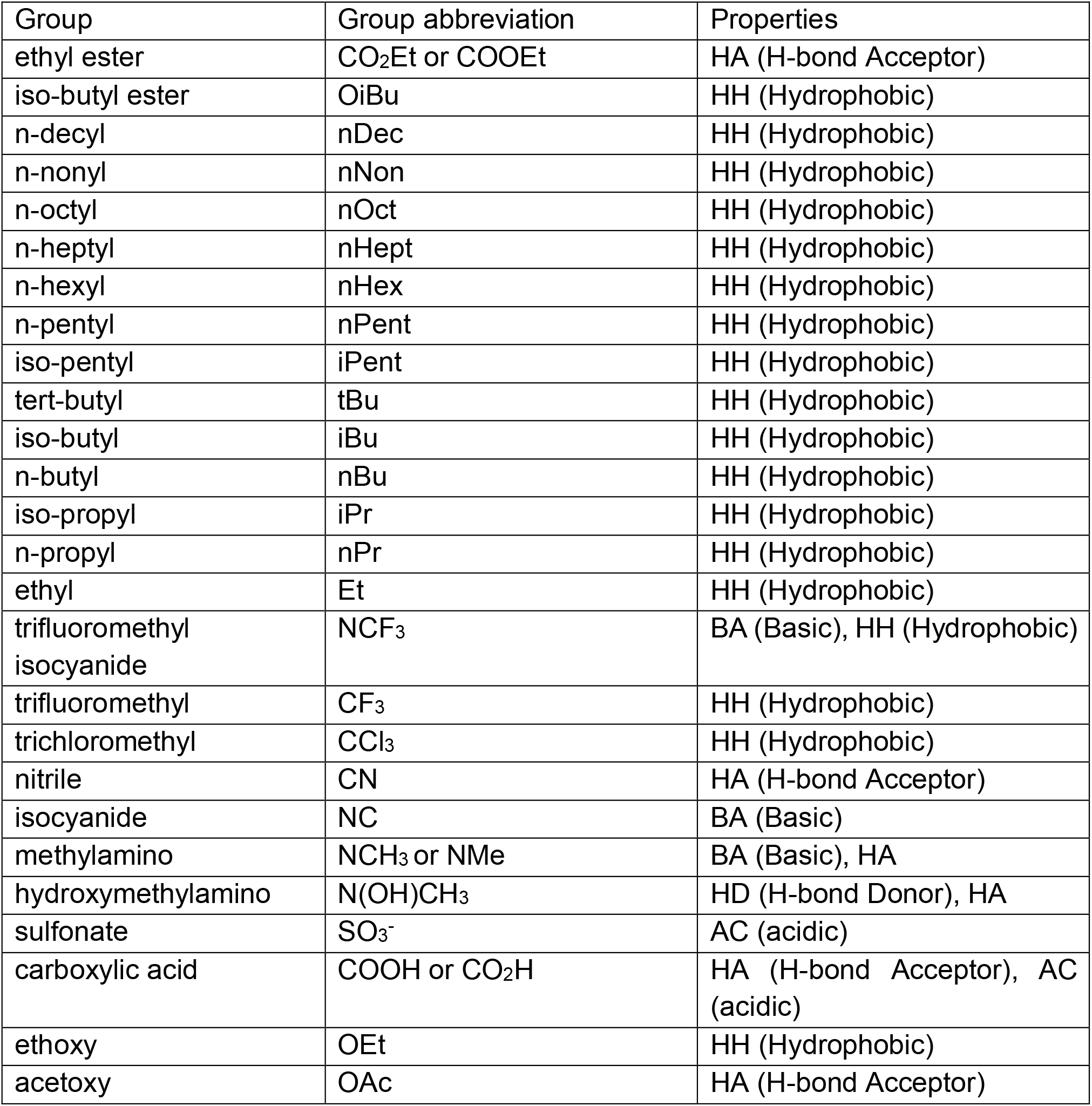
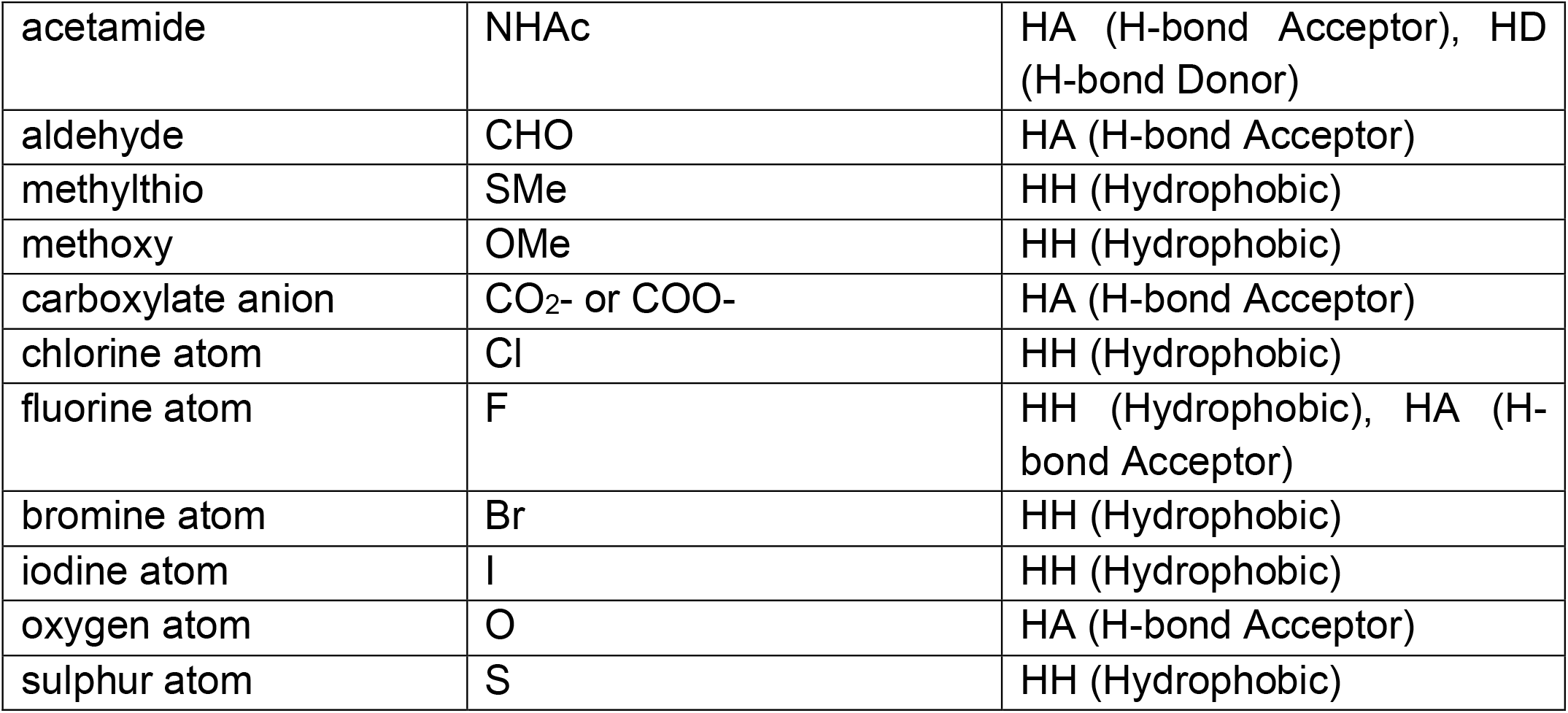
Functional groups and their properties categorized by CCGM metrics.

#### Chemotype and Pharmacophore Similarity Calculation

Compound similarities in CCGM are quantified using two distinct Dice similarity scoring functions: *chemotype similarity*, which assesses the structural similarity of compound graphs, and *pharmacophore similarity*, which evaluates the similarity of pharmacophore features. The final CCGM similarity score is calculated as a weighted average of these two metrics, with equal weights (0.5) applied by default. However, users can adjust this weight based on the specific requirements of their screening tasks. By default, CCGM assigns a weight of 1 to all nodes and edges in the template graph, an approach that is especially effective when screening large compound libraries, where the objective is to filter for compounds with similar chemotypes. When the core scaffold and R-groups are identified, users can adjust the node and edge weights for the scaffold of interest, leading to more refined similarity search results as shown in Fig. 1C. In addition to comparing a graphs’ nodes and edges, CCGM incorporates pharmacophore similarity by comparing pharmacophore features between the template and query molecules, as described previously. Together, these distinct similarity scores ensures that CCGM identifies and filters compounds that not only share structural similarity but also exhibit comparable pharmacophore features.

#### Negative Weights in Chemotype Optimization

Exclusion Volume/Area is a key concept in quantitative structure-activity relationship (QSAR) studies, identifying specific R-groups or molecular regions where exploration could lead to unfavorable steric clashes or loss of selectivity. Current 2D representations and their associated screening approaches often fail to account for exclusion areas. CCGM addresses this limitation by introducing negative weights for specific nodes in the graph to penalize unfavorable molecular modifications. When extending an exit vector (R-group) in a particular direction is likely to compromise a compound’s potential as a lead candidate, users can assign a dummy atom (denoted as *) to represent exclusion areas. Negative weights can then be applied to the corresponding nodes and edges, as illustrated in Fig. 2B. By annotating template molecule in this manner, CCGM can exclude such molecules during screening or assign them a low CCGM similarity score, ensuring they are rank lower. This approach enables a more targeted and efficient compound selection process, prioritizing molecules with higher potential for success.

## Results & Discussion

CCGM utilizes a graph-based framework to represent core chemotype and pharmacophore features, enabling efficient and accurate identification of structurally similar compounds using distinct similarity metrics. To assess the performance of the CCGM, we conducted three different benchmarking experiments to assess the accuracy, reliability, and usability of CCGM compared to other compound similarity estimation methods. These benchmarks focused on evaluating CCGM, weighted CCGM (wCCGM), and other similarity metrics, such as Tanimoto and DeCAF, across various chemotypes and compound libraries. First, we calculated hit rates by identifying similar chemotypes. Next, we extended the evaluation to diverse chemotypes to benchmark CCGM’s robustness. Finally, to further demonstrate CCGM’s broad applicability, we applied the model to screen large commercial chemical and guide generative designs.

### Benchmarking across similar chemotypes

To evaluate the effectiveness of CCGM and wCCGM in identifying structurally and functionally similar compounds, we selected four FDA-approved drugs where the core chemotypes are similar, which includes compounds gefitinib, afatinib, neratinib, and dacomitinib (Fig. 3A). Specifically, all the four compounds contain a 4-quinazolinamine core scaffold, serving as the basis for identifying structurally similar molecules (Fig. 3B). To prepare a benchmark set, we retrieved similar compounds for each of the four FDA-approved drugs from the MolPort screening library (www.molport.com). This process resulted in a curated collection of 95 compounds sharing the same chemotype. Using each drug as a template, we applied CCGM and wCCGM, alongside a fingerprint-based method (Tanimoto similarity) and pharmacophore-based method (DeCAF), to virtually screen the dataset for potential hits. A similarity threshold of 0.75 was set to define a hit, enabling a direct comparison of each method’s effectiveness in identifying structurally analogous compounds to respective templates.

**Figure 3.**
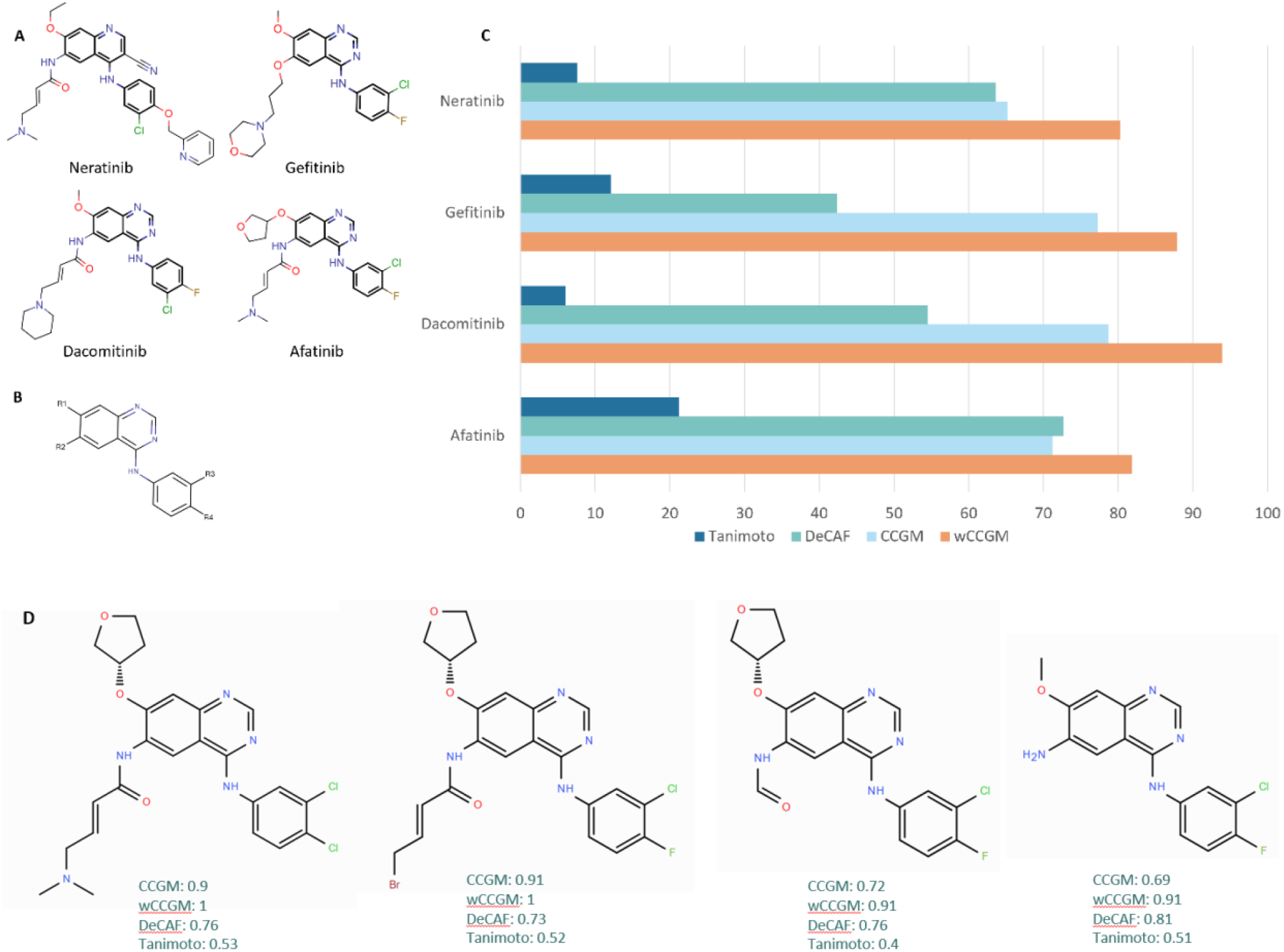
(A) Analogues of FDA-approved drugs sharing the same chemotype were selected: gefitinib, neratinib, dacomitinib, afatinib, and used as templates. A compound collection was prepared combining similar compounds for the four compounds from MolPort database, resulting in a dataset of 95 compounds (B) The core scaffold commonly shared among the template drugs (C) A bar plot showing the hit rates, at a similarity threshold of 0.75, calculated using different similarity functions (Tanimoto, DeCAF and CCGM and weighted CCGM) for template molecule to filter compounds from the dataset of similar chemotypes. (D) The figure represents the impact of minor structural modifications and solvent changes on various similarity scores similarity metrics (CCGM, wCCGM, Tanimoto and DeCAF) on compounds with similar chemotypes as dacomitinib.

The results of this benchmarking study, summarized in Fig. 3C, demonstrate that both CCGM and wCCGM reliably identified structurally similar compounds, with hit rates comparable to or exceeding those of traditional fingerprint-based and pharmacophore-based methods. Notably, wCCGM showed enhanced sensitivity in detecting subtle chemotype variations within the dataset, highlighting its potential as a robust tool for virtual screening and lead optimization.

### Benchmarking across diverse chemotypes

To further validate the reproducibility and robustness of CCGM, we applied it to a range of diverse chemotypes, specifically Type-1 kinase inhibitors from FDA-approved drug collection. Following the approach described earlier, the benchmark datasets were generated by retrieving structurally similar compounds from the MolPort database, excluding any drugs with fewer than 15 similar compounds. To ensure thorough evaluation, a comprehensive dataset was prepared by combining each of the benchmark set with 1000 random compounds (decoys). We then applied CCGM, wCCGM, DeCAF, and Tanimoto similarity metrics to screen this dataset, measuring hits at 10% enrichment factor. The metric quantifies the fraction of active compounds present in the top 10% of the ranked list, offering insight into each method’s screening efficiency. As shown in Fig. 4A, wCCGM demonstrated consistent performance across all chemotypes, gaining a higher hit rate compared to Tanimoto similarity and performance comparable to DeCAF similarity. These results underscore wCCGM’s enhanced ability to capture subtle variations in chemotype structure and pharmacophore features, illustrating its robustness and precision in virtual screening across structurally diverse compound classes.

**Figure 4.**
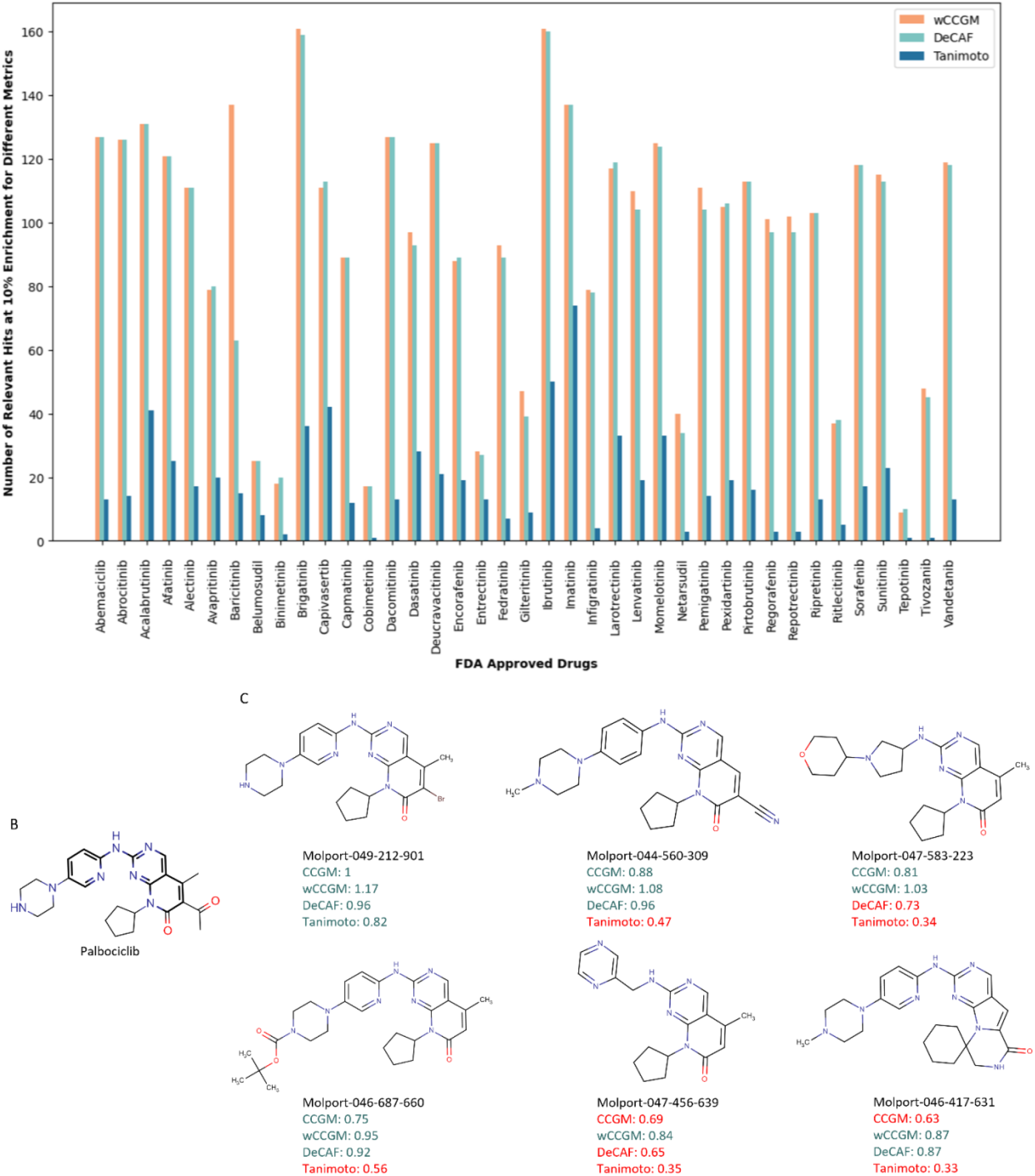
(A) Comparison of similarity scores across different metrics (wCCGM, Tanimoto, and DeCAF) based on the number of relevant hits at 10% enrichment factor at a similarity threshold of 0.75. (B) Structure of palbociclib. (C) Performance analysis of CCGM and wCCGM against Tanimoto, DeCAF similarity metrics on similar chemotype compounds of Palbociclib: Impact of minor structural modifications and solvent changes on similarity scores.

To provide deeper insight into the model’s performance, we analyzed palbociclib, a selective inhibitor of cyclin-dependent kinases (CDKs), as a representative template (Fig. 4B). Compounds with minor modifications, such as Molport-049-212-901 (with a bromine substitution), Molport-044-560-309 (bearing cyano and methyl groups), and Molport-047-583-223 (modified solvent-exposed region), displayed high similarity scores across all metrics, including CCGM, wCCGM and DeCAF (Fig. 4C). However, as structural changes grew more substantial, for example, in Molport-046-687-660 (with a bulkier solvent region), Molport-047-456-639 (with a distinct solvent environment), and Molport-046-417-631 (modified front-pocket region), the performance of Tanimoto similarity declined, indicating its limited sensitivity to these types of structural variations (Fig. 4C). Similarly, modifications in the solvent-exposed regions reduced DeCAF performance, likely due its reliance on rigid pharmacophore definitions that are less responsive to subtle changes. In contrast, wCCGM maintained consistently high performance despite increasing structural modifications, highlighting its robustness in accommodating diverse chemotypes and variations across different molecular regions.

### CCGM for screening large libraries

CCGM can effectively filter compounds with similar chemotype from large databases and can also guide the generative model to design molecules without deviating from a desired template. To illustrate these applications, we screened large compound libraries, such as MolPort (>500k compounds), using gefitinib as a template. This screening identified several promising analogs and molecules with high similarity scores to the gefitinib (Fig. 3A), confirming that they share the same chemotype. As shown in Fig. 5A, close analogs such as Molport-047-152-632, Molport-047-482-767, and Molport-047-950-123, were successfully identified. Additionally, we uncovered several structurally divergent molecules with low CCGM scores, examples include molecules such as Molport-010-672-516, Molport-000-087-168, and Molport-000-087-176. These molecules contain modifications, such as substitutions at positions distinct from the template chemotype or alterations in the solvent-exposed and front-pocket regions, contributing to their reduced similarity to gefitinib. This highlights CCGM’s utility in identifying structurally similar chemotypes within large compound libraries, while also revealing compounds with structural variations that can inform further exploration and optimization.

**Figure 5.**
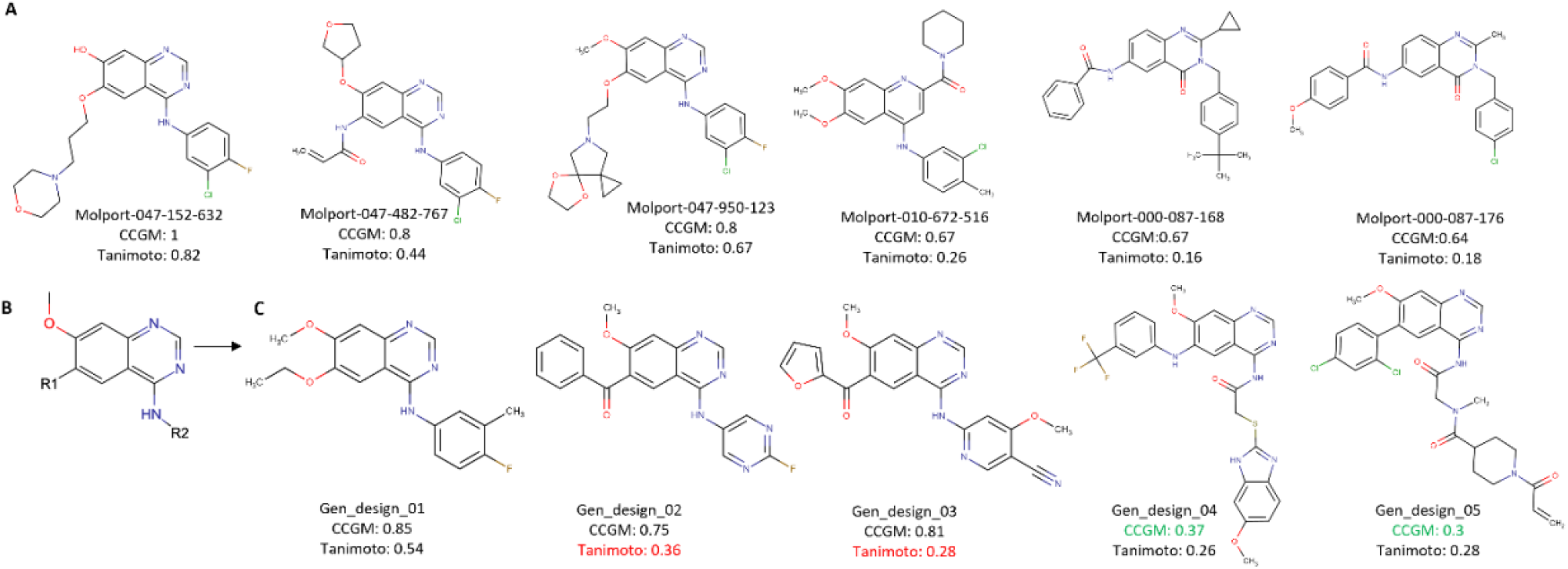
(A) Examples of compounds following virtually screening MolPort database using gefitinib as the template molecule, their respective CCGM similarity scores and Tanimoto similarity scores are given below hit compound. (B) The core scaffold of gefitinib used in as input for the in-house generative model. (C) A sample of CCGM-guided generative task where a series of gefitinib-like molecules were generated and their similarity with gefitinib was estimated using different similarity metrics.

Furthermore, CCGM can inform generative models to design new molecules that retain essential characteristics of the template. For instance, we used the gefitinib template with exit vectors defined as R1 and R2 in Fig. 5B. We steered an in-house generative model to produce gefitinib-like structures. The CCGM framework provides feedback when a generated molecule deviates from the core template structure. As illustrated in Fig. 5C, molecules generated by the model that closely align with the template (e.g., Gen_design_01, Gen_design_02, Gen_design_03), achieve high CCGM scores. In contrast, generated molecules that diverge from the template (e.g., Gen_design_04, Gen_design_05), exhibit low CCGM scores, highlighting the structural deviations.

The overall results emphasize the strength of CCGM and wCCGM in virtual screening tasks. Whether applied to a single template or a compound set with diverse chemotypes, CCGM consistently outperformed traditional fingerprint-based methods and pharmacophore-based screening approaches. By integrating both chemotype and pharmacophore features and offering the flexibility of weighted similarity metrics, wCCGM consistently achieved better results, making it a powerful and versatile tool for early-stage drug discovery.

## Conclusion

CCGM is an open-source Python framework specifically designed to support cheminformaticians during the hit-to-lead optimization phase of drug discovery. The framework employs a reduced graph representation of molecules to estimate molecular similarities. The CCGM similarity score integrates both chemotype and pharmacophore similarity metrics, enabling users to filter for compounds that share similar topology and pharmacophore characteristics. We demonstrated the utility of CCGM in screening and identifying hits across both similar and diverse chemotypes. In these tests, CCGM achieved accuracy levels comparable to or exceeding those of traditional fingerprint methods, such as Tanimoto, and pharmacophore-based approaches like DeCAF. Additionally, CCGM can serve as a virtual screening filter for large compound libraries and guide generative models, effectively isolating compounds with desired properties. Beyond conventional representations and scoring functions, we believe CCGM will be an invaluable addition to a medicinal chemist’s toolkit, accelerating and guiding the design and optimization of hits.

## Availability of data and materials

The dataset supporting the conclusions of the article are included within the article and its additional files.

## Authors’ contributions

Conceptualization: M.P, B.R, R.R; methodology: N.S, M.P, B.R; implementation:N.S, B.R; writing: N.S, B.R; reviewing: M.P, R.R.

## References

1. Hu Y, Stumpfe D, Bajorath J (2016) Computational Exploration of Molecular Scaffolds in Medicinal Chemistry. 4062–76. 10.1021/acs.jmedchem.5b01746

2. Nguyen-Vo T-H, Teesdale-Spittle P, Harvey JE, Nguyen BP (2024) Molecular representations in bio-cheminformatics. Memetic Comp 16:519–536. 10.1007/s12293-024-00414-6

3. An X, Chen X, Yi D, et al (2022) Representation of molecules for drug response prediction. Briefings in Bioinformatics 23:bbab393. 10.1093/bib/bbab393

4. Kearnes S, McCloskey K, Berndl M, et al (2016) Molecular graph convolutions: moving beyond fingerprints. J Comput Aided Mol Des 595–608. 10.1007/s10822-016-9938-8

5. Rogers D, Hahn M (2010) Extended-Connectivity Fingerprints. J Chem Inf Model 50:742–754. 10.1021/ci100050t

6. Bender BJ, Gahbauer S, Luttens A, et al (2021) A practical guide to large-scale docking. Nat Protoc 16:4799–4832. 10.1038/s41596-021-00597-z

7. Hawkins P, Skillman AG, Nicholls A (2007) Comparison of Shape-Matching and Docking as Virtual Screening Tools. Journal of Medicinal Chemistry 50:74–82. 10.1021/jm0603365

8. Stepniewska-Dziubinka M, Zielenkiewicz P, Siedlecki P (2017) DeCAF— Discrimination, Comparison, Alignment Tool for 2D PHarmacophores. Molecules 1128. 10.3390/molecules22071128

9. Rasha Atwi, Ye Wang, Simone Sciabola, Adam Antoszewski (2024) ROSHAMBO: Open-Source Molecular Alignment and 3D Similarity Scoring. Journal of Chemical Information and Modeling

10. Aric A. Hagberg, Daniel A. Schult, Pieter J. Swart Exploring network structure, dynamics, and function using networkX. In: Proceedings of the 7th Python in Science Conference (SciPy 2008). pp 11–15

11. Karulin B, Kozhevnikov M (2011) Ketcher: web-based chemical structure editor. Journal of Cheminformatics 3:P3. 10.1186/1758-2946-3-S1-P3

12. Weininger D (1988) SMILES, a chemical language and information system. 1. Introduction to methodology and encoding rules. J Chem Inf Comput Sci 28:31–36. 10.1021/ci00057a005

13. Dalke A, Hastings J (2013) FMCS: a novel algorithm for the multiple MCS problem. Journal of Cheminformatics 5:O6. 10.1186/1758-2946-5-S1-O6

